# Maramycin, a cytotoxic isoquinolinequinone terpenoid produced through heterologous expression of a bifunctional indole prenyltransferase /tryptophan indole-lyase in *S. albidoflavus*

**DOI:** 10.1101/2024.02.14.580309

**Authors:** Matiss Maleckis, Mario Wibowo, Sam E. Williams, Charlotte H. Gotfredsen, Renata Sigrist, Luciano D.O. Souza, Michael S. Cowled, Pep Charusanti, Tetiana Gren, Subhasish Saha, José M. A. Moreira, Tilmann Weber, Ling Ding

## Abstract

Isoquinolinequinones represent an important family of natural alkaloids with profound biological activities. Heterologous expression of a rare bifunctional indole prenyltransferase /tryptophan indole-lyase enzyme from *Streptomyces mirabilis* P8-A2 in *S. albidoflavus* J1074 led to the activation of a putative isoquinolinequinone biosynthetic gene cluster and production of a novel isoquinolinequinone alkaloid, named maramycin (**1**). The structure of maramycin was determined by analysis of spectroscopic (1D/2D NMR) and MS spectrometric data. The prevalence of this bifunctional biosynthetic enzyme was explored and found to be a recent evolutionary event with only a few representatives in Nature. Maramycin exhibited moderate cytotoxicity against human prostate cancer cell lines, LNCaP and C4-2B. The discovery of maramycin (**1**) enriched the chemical diversity of natural isoquinolinequinones and also provided new insights into crosstalk between the host biosynthetic genes and the heterologous biosynthetic genes in generating new chemical scaffolds.

## Introduction

*Streptomyces* represent a prolific source for bioactive secondary metabolites, exemplified by the antibiotics streptomycin, tetracycline and daptomycin, and the anticancer drugs doxorubicin and bleomycin, all WHO essential medicines^1^.

The advances of modern genome mining analyses have revealed that still a large fraction of the biosynthetic potential is untapped: 70% of the secondary metabolites produced by *Streptomyces* are cryptic compounds whose corresponding genes are normally silent in standard laboratory culture conditions^2–5^. Many approaches, especially synthetic biology and ecology have been applied to access this genetic potential^6,7^ by using, for example, genetic engineering^8^, heterologous expression of a BGC in another host^9^, chemical elicitors^10^ and co-cultivation^11^. These strategies are often employed to unravel the potential of metabolite production of *Streptomyces* through cryptic gene activation, allowing for a vast number of potentially valuable compounds to be discovered.

Isoquinolinequinones represent an important family of natural alkaloids with profound biological activities. They are predominantly isolated from marine invertebrates, such as cytotoxic caulibugulones and perfragilins from the bryozoan *Caulibugula inermis*^12^ and *Membranipora perfragilis*^13^, antineoplastic cribrostatins from the sponge *Cribrochalina* sp.^14^ and antimicrobial and anti-inflammatory renierones from the sponge *Renier* sp.^15^ and *Haliclona* sp.^16^. Hence, it was believed that isoquinolinequinones are natural products from marine invertebrates. However, the recent discovery of such compounds including the mansouramycins^17–20^ and albumycin^21^ from *Streptomyces* showed that microbes are also isoquinolinequinones producers. An isoquinolinequinone biosynthetic gene cluster in *S. albidoflavus* J1074 (previously recognized under the name *S. albus* J1074) was proposed by Chai et al.^22^ where they identified three bicyclic isoquinolinequinone products when expressing a putative BGC in *S. coelicolor* M1146^19^. This BGC was further studied by Shuai et al.^23^, who confirmed mansouramycin production in *S. albidoflavus* Del14^24^, a genome minimized strain of *S. albidoflavus* J1074^25^. Through feeding studies and NMR analysis, tryptophan was identified to be a precursor in the biosynthesis of mansouramycin D and tryptophan derived intermediates were detected in knockout strains^23^.

*S. albidoflavus* J1074 has been routinely used as a heterologous expression host system in our laboratory^26–28^. Although it harbors a putative mansouramycin BGC, we have not detected production of any related alkaloids under the laboratory conditions we routinely use. In a recent study, during expression of an azodyrecin BGC into *S. albidoflavus* J1074^29^, we serendipitously noticed the production of an unknown alkaloid (**1**). In this study, we link the production of this novel alkaloid, which we named maramycin (**1**), belonging to the isoquinolinequinone family, to the heterologous expression of a putative-bifunctional indole prenyltransferase /tryptophan indole-lyase. Furthermore, we describe isolation, structure elucidation and biological activities of maramycin (**1**).

## Results and Discussion

### Production of maramycin, a novel isoquinolinequinone terpenoid through heterologous expression of *mara1*

In our recent study on the biosynthesis of azodyrecins^29^, we heterologously expressed a putative *azd* BGC for cluster validation. Untargeted metabolomic analysis of LC-MS data revealed the production of a novel metabolite (**1**) that was not detected in the control strain not carrying the BGC (**Figure S1**). Detailed analysis of HRESIMS data revealed a molecular formula C21H19N3O2 for **1**, with a double bond equivalent (DBE) of 14. Analysis of its UV spectrum and MS/MS fragmentation pattern (**Figure S2**) suggested that **1** was unrelated to the biosynthesis of the azodyrecins (DBE = 3). It was unclear if an induced production of azodyrecin in the heterologous host has activated cryptic BGC in *S. albidoflavus* J1074^22^ or there was interplay between native and heterologous enzymes resulting in the production of **1**.

To elucidate the structure of the novel alkaloid **1**, *S. albidoflavus* J1074 bearing the *azd* BGC was cultivated on 5 L of solid agar and extracted using ethyl acetate. Following various isolation and purification steps, **1** was obtained and characterized by detailed NMR (**Table S1)** and MS analysis. Compound **1** was isolated as a red amorphous powder. The 1D ^13^C (**Figure S3**), 2D^1^H-^13^C edited-HSQC and HMBC (**Figure S4**) spectra identified 21 carbon signals, including three methyl groups, a methylene, six methine groups, and eleven quartenary carbons. The ^1^H NMR spectrum (**Figure S5**) of **1** showed signals from three methyl groups [δH 1.75 (6H) and 2.84], one methylene appearing as a doublet, six sp^2^ methines, and two exchangeable protons belonging to NH groups (δH 7.86 and 11.91); the latter two signals were confirmed by HSQC and ^1^H-^15^N HMBC (**Figure S6**) experiments. These data indicated that compound **1** belonged to the isoquinolinonequinone class of compounds. Furthermore, the methine proton resonances appearing as singlets at δH 8.97 and 5.69, together with the signals of NH at δH 7.86 and the methyl doublet at δH 2.84 were characteristics of mansouramycins^18^, a group of isoquinolinonequinones previously isolated from a marine *Streptomyces*^18^. Following literature reviews and spectroscopic data comparison, the NMR data of **1** were found to be similar to those of mansouramycin E^18^, except for the presence of an additional prenyl unit in **1**, and that an aromatic methine in mansouramycin E was present as a quartenary carbon (C-3′) in **1**. The presence and position of an additional prenyl group was established following 2D NMR data analysis (**Figure 1**); for instance, COSY (**Figure S7**) correlations between CH2-6′ and CH-7′ together with HMBC correlations from both H-9′ and H-10′ to C-8′ and C-7′. Further HMBC correlation from H-7′ to C-3′ positioned the prenyl moiety on C-3′. The remaining structure of **1** was further confirmed by detailed HMBC data (**Figure S8**) analysis. Hence, the structure of **1** was established as a new isoquinolinequinone terpenoid and named maramycin (**1**).

**Figure 1.**
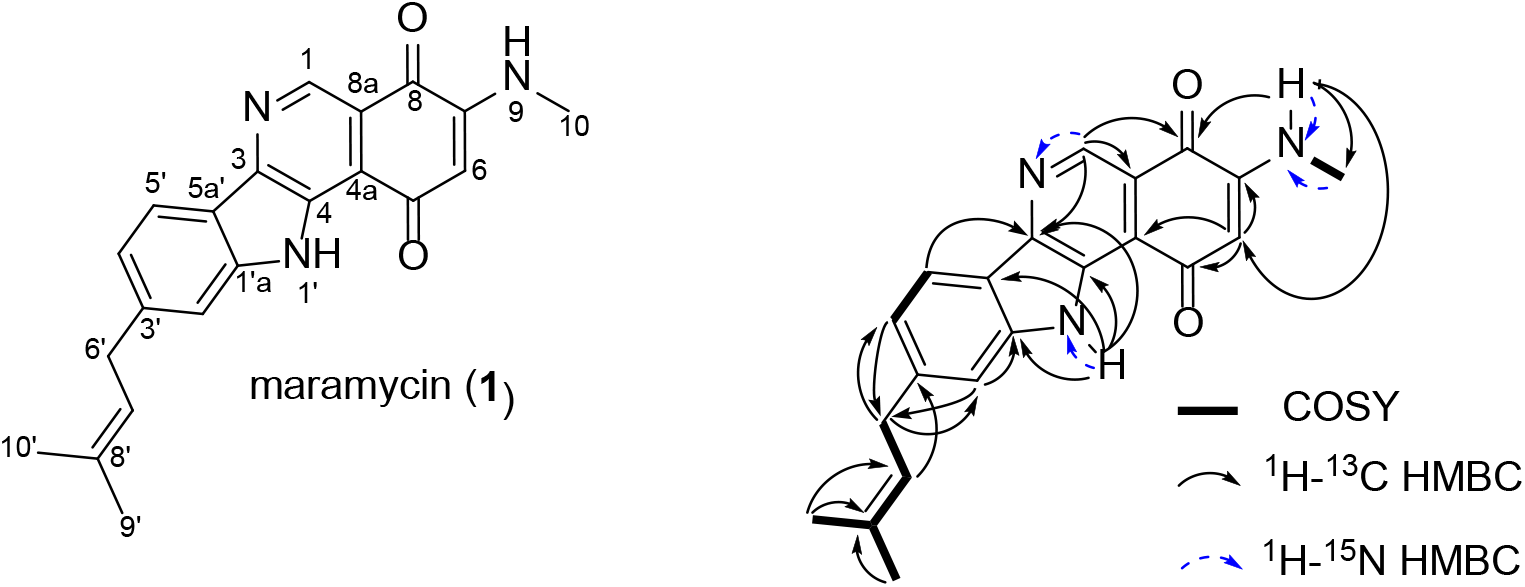
Chemical structure of isoquinolinequinone terpenoid maramycin (**1)** and selected 2D NMR correlations important for the structure elucidation.

### Biosynthesis of maramycin

Mansouramycins are encoded by *man* BGC^23^ in *S. albidoflavus* J1074, which has been characterized through gene inactivation and heterologous expression experiments, showing that *mans3-9* are essential genes for mansouramycin biosynthesis^23^. There have been no previous reports on natural prenylated mansouramycins and there are no adjacent prenyltransferase enzymes in *S. albidoflavus* J1074 associated with the *man* BGC. Given the distinctive prenyl group present in **1**, we reexamined the cloned *azd* BGC region and identified a candidate gene (*mara1*). The putative Mara1 enzyme contained an indole prenyltransferase domain^30^ and a tryptophan indole-lyase domain^31^. To confirm its function, expression of just *mara1* in *S. albidoflavus* J1074 indeed resulted in the production of maramycin (**1**) (**Figure 2.A**).

**Figure 2.**
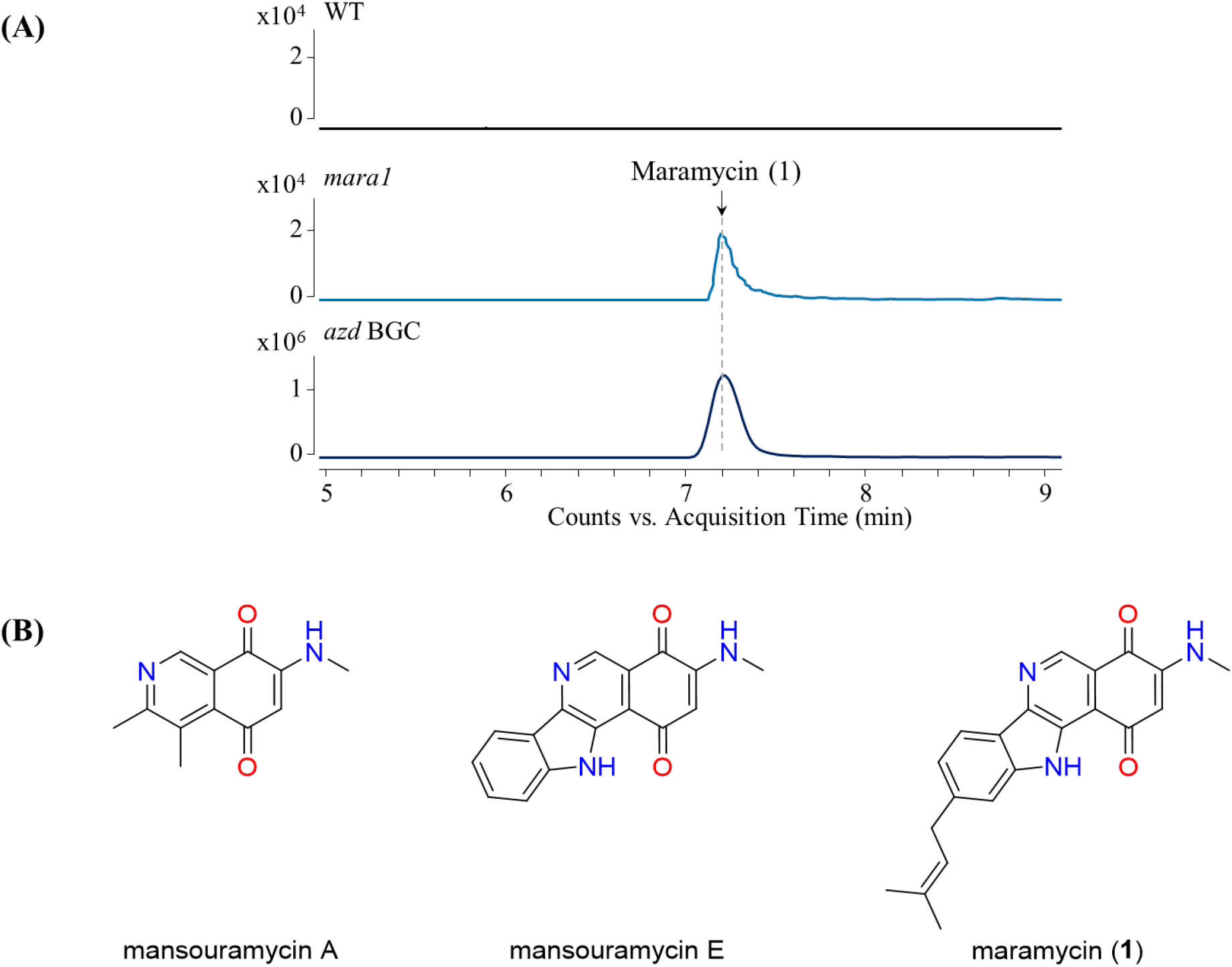
(**A**) Extracted ion chromatogram ([M+H]^+^ = 346.1550 ± 10 ppm) of maramycin (**1**) from LC-MS samples of *Streptomyces albidoflavus* J1074 (**WT**) and two mutant strain samples expressing *mara1* and *azd* BGC, respectively. (**B**) Structures of mansouramycins A^32^ and E ^22^, and maramycin (**1**).

Due to the high structural similarity to the mansouramycins, we hypothesize that maramycin was synthesized in combination of the native *man* BGC and the foreign *mara1* gene. The *man* BGC produces **2**, an intermediate for mansouramycin which we believe reacts with the prenylated indole product of Mara1 (**Figure 3**). Mara1 is a bifunctional enzyme which could catalyze the conversion of tryptophan into a prenylated indole. Firstly, a tryptophan is prenylated at the C-3′ position by the indole prenyltransferase, subsequently the tryptophan indole-lyase domain facilitates a β-elimination reaction producing the prenylated indole precursor 6-(3-methyl-2-butenyl)-indole. This prenylated indole is then incorporated into the quinone intermediate to form maramycin (**1**).

**Figure 3.**
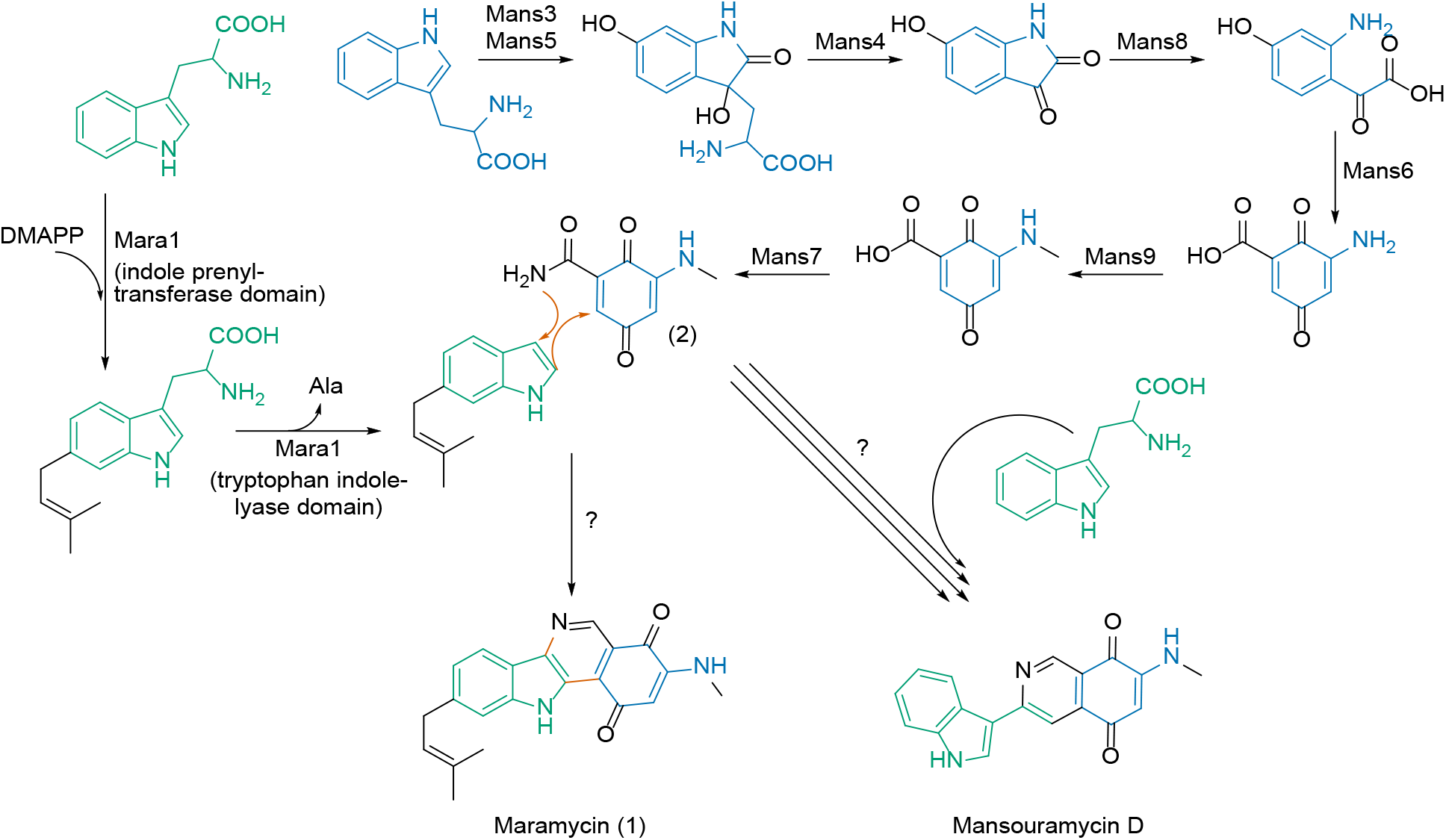
Proposed biosynthesis pathway of maramycin (**1**), where Mara1 is responsible for formation of 6-(3-methyl-2-butenyl)-indole which reacts with mansouramycin biosynthesis intermediate, 5-(methylamino)-3,6-dioxocyclohexa-1,4-diene-1-carboxamide (**2**), formed by Mans7 to form maramycin (**1**).

### Mara1: A rare bifunctional enzyme

The Mara1 enzymatic structure was assessed by comparing an AlphaFold protein model^33,34^ to crystal structures available in the Protein Data Bank (PDB) using FoldSeek^35^. This analysis confirmed the similarity of the Mara1 N-terminus to the prenyltransferase, PriB, (PDB: 5INJ, 898 MatchAlign score, 0.923 Root Mean Square Deviation “RMSD”)^36^ and the C-terminus to Tryptophan indole-lyase (PDB: 5W1B, 1672 MatchAlign score, 0.745 RMSD)^37^. These alignment scores demonstrate high structural congruence between the two distinct domains and the fused Mara1 enzyme (**Figure 4.A**).

**Figure 4.**
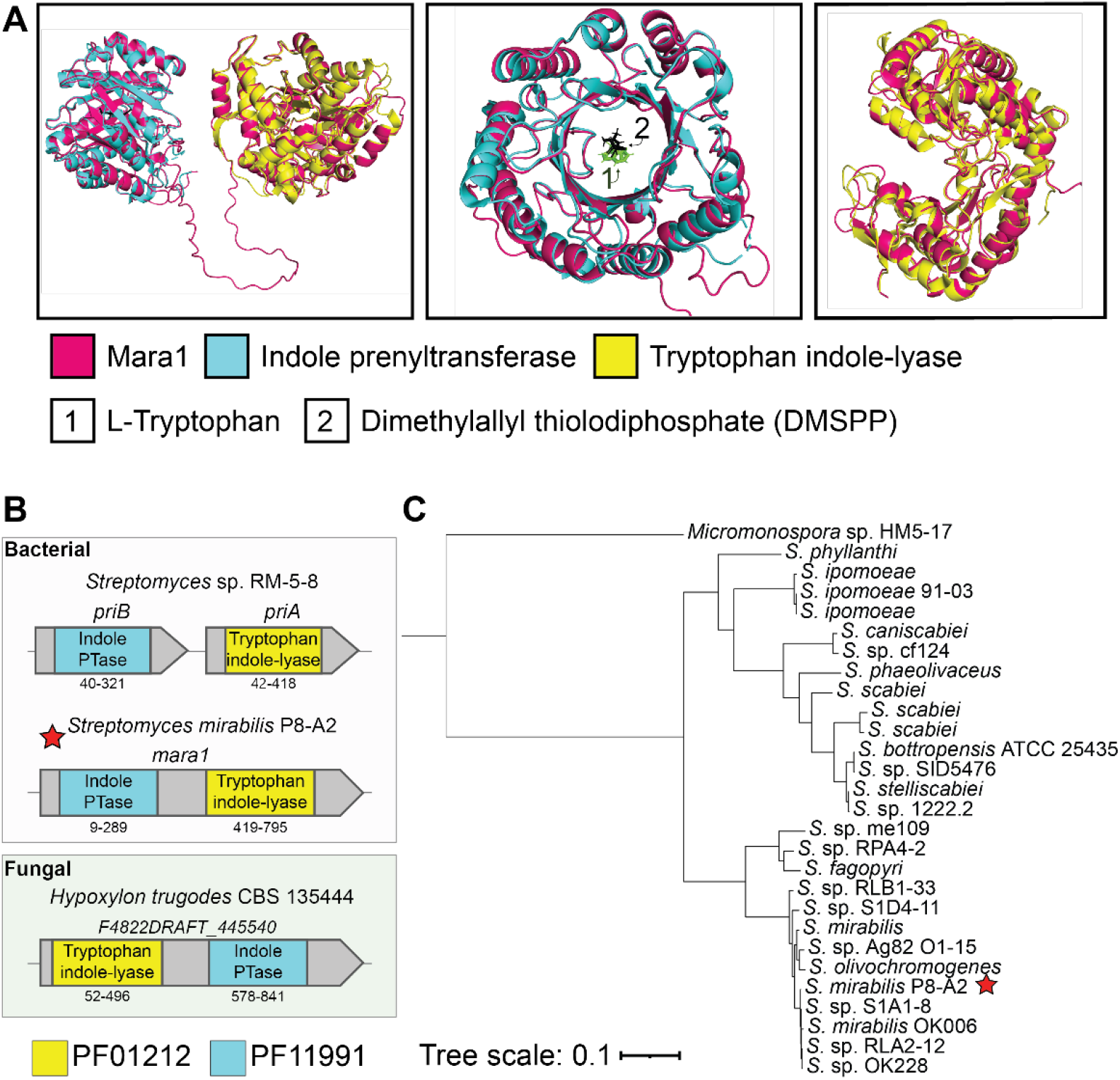
**(A)** Superimposition of Mara1 AlphaFold model to indole prenyltransferase and tryptophan indole-lyase records deposited in Protein Data Bank (PDB). The prenyltransferase, PriB crystal structure was made while in ternary complex with L-tryptophan and dimethylallyl thiolodiphosphate (DMSPP). **(B)** Pfam domain architecture present in *S. mirabilis* P8-A2 *mara1* in comparison to the separate PriB and PriA genes involved in the production of 5-isoprenylindole-3-carboxylate β-D-glycosyl ester from *Streptomyces* sp. RM-5-8. A reversed domain architecture is seen in fungi. The amino acid position of the identified domain is listed. **(C)** A phylogenetic tree of the bacterial homologues of the bifunctional indole enzyme. The red star on each panel highlights *S. mirabilis* P8-A2.

To investigate the prevalence of other proteins with similar fused domain architecture we queried the Interpro database (v 96.0) for proteins containing the Pfam domains 11991 (indole-prenyltransferase) and Pfam 01212 (tryptophan indole-lyase). Our query produced two distinct families of proteins: one bacterial (29 hits) and one fungal (30 hits). The bacterial family was almost exclusively found in *Streptomyces*, apart from a single homologue found in the taxonomically distinct but related genus *Micromonospora*. These proteins featured a N-terminal indole prenyltransferase domain followed by C-terminal tryptophan indole-lyase domain. Interestingly, the fungal enzymes displayed a reversed domain order, with the tryptophan indole-lyase at the N-terminus (**Figure 4.B**). Analysis using the antiSMASH database (v4)^38^ identified the bacterial bifunctional enzyme in 28 biosynthetic regions (*Micromonospora* sp. HM5-17 not present antiSMASH database v4). Further 156 regions contained the two domains as separate, adjacent enzymes, all of which maintained the sequential domain order seen in the Mara1 bifunctional enzyme. This separate gene architecture is also observed in the PriB containing BGC responsible for 5-isoprenylindole-3-carboxylate β-D-glycosyl ester production^39^ (MiBIG: BGC0001483) and the isatin-type antibiotic 7-prenylisatin^40^ (MiBIG: BGC0001294) (**Figure 4.B**). In both BGCs these domains act together to produce a prenylated indole structure from a tryptophan precursor. *Streptomyces* strains containing this enzyme are closely related, suggesting a potential recent gene fusion event producing this novel Mara1 enzyme (**Figure 4.C**).

### Cytotoxicity of maramycin

Isoquinolinequinones have been reported to be potential anticancer drug candidates ^41^. The effects of maramycin (**1**) on cell viability and proliferation were assessed using various prostate cancer cell lines. Exposure to maramycin (10 μM) for 72 h, inhibited cell growth of LNCaP and C4-2B cells, by 77.9% and 64.1%, respectively (**Figure S9.A and S9.B, respectively**). This was caused by moderate cytotoxicity elicited by exposure to **1** which showed IC50 values of 11.8 μM and 18.4 μM, against LNCaP and C4-2B cells, respectively (**Figure S9.E**). Drug resistance remains the main limiting factor for drug efficacy in cancer treatment. Like for antimicrobial resistance, reduction of the intracellular concentration of a drug by enhancement of drug efflux from cells, is a key mechanism of resistance. Therefore, we evaluated the effect of maramycin on the multi-drug resistant sublines LNCaP^R^ and C4-2B^R^, which overexpress the ABCB1/P-glycoprotein (P-gp) efflux pump and found it to exert comparable levels of cell growth inhibition in these cells (**Figures S8.C and S8.D**), suggesting it can evade Pgp-mediated multidrug resistance. Maramycin does not appear to have antibacterial properties against drug resistant Gram-negative pathogens. When *S. albidoflavus* J1074 WT and *S. albidoflavus* J1074 bearing the *azd* BGC were co-cultured against multidrug-resistant strains of *Actinetobacter baumannii* and *Escherichia coli*, no inhibition zone was observed.

### Conclusion

In this study we discovered and characterized maramycin, a novel alkaloid with promising anticancer activity. The production of this compound arose from cross-talk of a rare bifunctional prenyltransferase/tryptophan indole lyase, capable of generating prenylated indoles from a tryptophan, and native mansouramycin biosynthesis. We believe this enzyme has wide applicability to rationally engineer tryptophan derived natural products and to create promising novel analogues.

## Methods

### General Experimental Procedures

Optical rotations were recorded on an AUTOPOL III - S2 Dual Wavelength (589/546 nm) Automatic Polarimeter (Rudolph Research Analytical). IR data were acquired on Bruker Alpha FTIR spectrometer using OPUS version 7.2. The NMR spectra were recorded on a Bruker AVANCE III 800 MHz spectrometer equipped with a 5 mm TCI CryoProbe using standard pulse sequences. The ^1^H and ^13^C NMR chemical shifts were referenced to the residual solvent signals at δH 2.50, δC 39.52 ppm for for DMSO-*d*6. UHPLC-HRMS was performed on an Agilent Infinity 1290 UHPLC system equipped with a diode array detector. UV−vis spectra were recorded from 190 to 640 nm. All solvents and chemicals used for HRMS, and chromatography were LC-MS grade, while the solvents for metabolite extraction were of HPLC grade. Water was purified using a Milli-Q system.

### LC-ESI-HRMS/MS Analysis

Ultra-high-performance liquid chromatography–diode array detection–quadrupole time-of-flight mass spectrometry (UHPLC–DAD–QTOFMS) was performed on an Agilent Infinity 1290 UHPLC system (Agilent Technologies, Santa Clara, CA, USA) equipped with a diode array detector. Separation was achieved on a 150 × 2.1 mm i.d., 1.9 μm, Poroshell 120 Phenyl Hexyl column (Agilent Technologies, Santa Clara, CA) held at 40°C. The sample (1 μL) was eluted at a flow rate of 0.35 mL min^−1^ using a linear gradient from 10% acetonitrile (LC-MS grade) in Milli-Q water buffered with 20 mM formic acid increasing to 100% in 10 min, staying there for 2 min before returning to 10% in 0.1 min. Starting conditions were held for 3 min before the following run.

Mass spectrometry (MS) detection was performed on an Agilent 6545 QTOF MS equipped with Agilent Dual Jet Stream electrospray ion source (ESI) with a drying gas temperature of 250°C, a gas flow of 8 L min^−1^, sheath gas temperature of 300°C and flow of 12 L min^−1^ Capillary voltage was set to 4000 V and nozzle voltage to 500 V in positive mode. Mass spectra were recorded as centroid data, at an *m/z* of 100–1700, and auto MS/HRMS fragmentation was performed at three collision energies (10, 20, and 40 eV), on the three most intense precursor peaks per cycle. The acquisition rate was 10 spectra s^−1^. Data were handled using Agilent MassHunter Qualitative Analysis software (Agilent Technologies, Santa Clara, CA). Lock mass solution in 70 % MeOH in water was infused in the second sprayer using an extra LC pump at a flow of 15 μL min^−1^ using a 1:100 splitter. The solution contained 1 μM tributylamine (Sigma-Aldrich) and 10 μM Hexakis (2, 2, 3, 3-tetrafluoropropoxy) phosphazene (Apollo Scientific Ltd., Cheshire, UK) as lock masses. The [M + H]^+^ ions (*m/z* 186.2216 and 922.0098, respectively) of both compounds were used.

### Microbial strains and culture conditions

*Escherichia coli* ET12567/pUZ8002^42,43^, *Escherichia coli* Mach1 (Thermo Fisher Scientific), *Streptomyces albidoflavus* J1074^25^ and *Streptomyces albidoflavus* J1074 ΦC31::pAzd^29^. All *Escherichia coli* strains were grown in liquid/solid LB medium (5.0 g/L yeast extract, 10.0 g/L peptone, 10.0 g/L NaCl) at 37 °C. *Streptomyces albidoflavus* was grown on SFM (20.0 g/L fat reduced soy flour (fettreduziert Bio Sojamehl; Hensel, Germany)), 20.0 g/L D-mannitol (Sigma-Aldrich), and 1.0 L tap water (Kgs. Lyngby, Denmark)) or ISP2 (yeast extract 4g/L (Thermo Fisher Scientific), malt extract 10g/L (Sigma-Aldrich), glucose 4g/L (Sigma-Aldrich), 1.0 L deionized water) at 30 °C. For conjugations, SFM media was supplemented to contain final concentration of 10 mM MgCl2. Appropriate antibiotics were supplemented with the following working concentrations: 100 μg/mL apramycin sulfate (Sigma-Aldrich), 25 μg/mL chloramphenicol (Sigma-Aldrich), 50 μg/mL kanamycin sulphate (Sigma-Aldrich) and 25 μg/mL nalidixic acid (Sigma-Aldrich).

### Heterologous expression of *mara1* in *S. albidoflavus* J1074

All polymerase chain reactions were performed using Q5 High-Fidelity 2X Master Mix (New England Biolabs) and all primers were synthesized by IDT (Integrated DNA Technologies). PCR amplification of pRM4e^44^ was performed using matmal0292: “GCGAGTGTCCGTTCGAG” and matmal0293: “ATGGACGTCCCCTTCCT”, while *mara1* was amplified by primer extension PCR using matmal0294: “actcgaacggacactcgcCTAGACGGTCACCGGCTG” and matmal0296: “caggaaggggacgtccatATGATCACCTCCCGTCCAGG”. The PCR products of expected 5.4 kbp and 2.6 kbp size, respectively, were gel purified using NucleoSpin Gel and PCR Clean-up (Macherey-Nagel) kit according to suppliers’ instructions. The *mara1* fragment was cloned into pRM4e vector using NEBuilder HiFi DNA Assembly (New England Biolabs) master mix and introduced into *E. coli* Mach1 by heat shock method^45,46^. The obtained plasmid, pRM4e-*mara1* (**Figure S10**), was verified via colony PCR, expected fragment size 3.1 kbp, and subsequent Sanger sequencing (Eurofins Genomics) using matmal0208: “GTCTGTCGAGAAGTTTCTGATC” and matmal0209: “ACATGTTCTTTCCTGCGTTATC”. The acquired plasmid was purified using NucleoSpin Plasmid EasyPure (Macherey-Nagel) kit according to suppliers instructions, cloned into *E. coli* ET12567/pUZ8002 and introduced into *S. albidoflavus* J1074 via conjugation. The resulting apramycin resistant clones were selected for LC-MS analysis.

### Bacterial cultivation

A seed culture was prepared by inoculating spores of strain *S. albidoflavus* J1074 ΦC31::pAzd into a baffled conical flask containing 50 mL of liquid ISP2 medium and incubated at 30 °C overnight with constant shaking at 180 rpm. The seed culture was inoculated on ISP2 agar plates and incubated at 30 °C for 7 days in the dark. A total of 5 L (200 plates) of ISP2 agar were prepared for extraction and isolation of maramycin.

### Extraction and isolation of maramycin

The agar cultures were sliced into small pieces and extracted under sonication for 30 min with EtOAc (2 × 5 L). The EtOAc was filtered and removed under reduced pressure using rotary evaporator to yield 599 mg of crude extract. The extract was subjected to flash chromatography using a C18-bonded Si-gel cartridge (Biotage SNAP 50 g) on Biotage Isolera Flash Chromatography system, eluted with step-gradient solvent systems (10% MeOH/H2O to 100% MeOH; 10% increment; 132 mL each) at a flow rate of 30 mL/min to give 20 fractions (66 mL each). Fractions 14–20 were combined following HRMS analysis that suggested the presence of the isoquinolinquinone terpenoid. The combined fraction was further purified using RP-HPLC column (Luna 5 μm C18-Phenomenex, 100 Å, 250 × 10 mm) with a linear gradient of 45%–85% ACN/H2O in 20 min to afford maramycin (**1**, 1.1 mg).

### Bioinformatic analysis of bifunctional enzyme

The *mara1* sequence is identical to already existing model AF-A0A856NK67-F1-model_v4 in the AlphaFold database^33,34^. Structurally similar models were identified in the PDB using FoldSeek^35^. The similarity of identified matches to Mara1 were assessed by performing PyMOL “super” structure alignment. The BGC annotation tool antiSMASH 7.0.0^47^was used to annotate the *mara1* gene and identify relevant Pfam domains. To identify further proteins containing both Pfam 01212 (β-eliminating lyase) and Pfam 11991 (tryptophan dimethylallyltransferase) domains, the InterPro database (v 96.0) was queried via the domain search. The 29 amino sequences of identified bacterial proteins were downloaded and aligned with MAFFT (v 7.490) using the --auto flag and model L-INS-I^48^. Phylogenetic trees were then constructed using FastTree2 (v 2.1.11)^49^and visualized and annotated in iTol (v 6)^50^. To identified enzymes in other biosynthetic gene clusters the amino acid sequence was queried against the antiSMASHdb (v4)^51.^.

### Cell growth and cytotoxicity assay

Two prostate cancer cell lines, LNCaP and C4-2B, as well as the drug-resistant derivative sublines, LNCaP^R^ and C4-2B^R^, respectively^52^, were used to evaluate the effect of maramycin on cell viability and proliferation. All cell lines were cultured and maintained in RPMI-1640 medium + glutaMAX™-I (Gibco, Invitrogen, Carlsbad, CA, United States) supplemented with 10% fetal bovine serum (FBS). One day prior to drug exposure measurements, cells were seeded into 6-well plates at a density of 0.3×10^6^ cells/well. On the next day, the medium in each well was replaced with 2mL of fresh warm medium containing either 10 μM of maramycin or vehicle. Cell proliferation dynamics were monitored in real-time using a lens-free Cellwatcher microscopy device (PHIO, Germany). The cell growth curves were generated with the analysis module available from PHIO to determine the total area covered by cells. Cytotoxicity was evaluated using the CellTox Green Cytotoxicity Assay kit (Promega, Madison, WI, USA) after 48h of exposure to various concentrations of maramycin (10^−7^ M to 2×10^−5^ M), according to manufacturer’s instructions. Briefly, CellTox green cytotoxicity reagent was added to the media at a final concentration of 1X, and the relative cytotoxicity was calculated relative to the control well treated with vehicle. The drug concentrations that caused inhibition of 50% cell viability (IC50) were determined from the dose-response curves. Curve fitting and analyses using non-linear regression models were performed using GraphPad Prism version 10.0.0 for Windows (GraphPad Software, Boston, MA, USA).

### *Maramycin* (1)

Red amorphous powder; UV (CH3CN/H2O) λmax 236 (100%), 270 (50%), 390 (25%) nm; IR (ATR) *v*max 3306, 2943, 2801, 2023, 1449, 1409, 1120, 1020, 623 cm^-1^; ^1^H and ^13^C NMR see Supporting Information (Table S1); (+)-HRESIMS *m/z* 346.1564 [M+H]+ (calcd for C21H20N3O2, 346.1550).

## Data availability

MassIVE MSV000093927, Genome sequence data for *Streptomyces mirabilis* P8-A2 is available under the NCBI RefSeq accession NZ_JARAKF000000000.1.

## Supporting Information

Additional details on the HPLC traces of heterologous expression strain, NMR data of maramycin, bioactivity data and plasmid map for heterologous expression of Mara1.

## Acknowledgements

This study was supported by the Danish National Research Foundation (DNRF137) as part of the Center for Microbial Secondary Metabolites (CeMiSt). S.E.W. would furthermore acknowledge funding by the Novo Nordisk Foundation Postdoctoral Fellowship (NNF22OC0079021). T.W. would furthermore acknowledge funding by the Novo Nordisk Foundation (NNF20CC0035580, NNF16OC0021746). The NMR Center DTU and the Villum Foundation are acknowledged for access to the 800 MHz spectrometer.

The metabolomic data was generated at DTU Metabolomics Core facilities with help of A. Andersen.

## Author contributions

M.M., M.W., S.E.W set out the methodology of the study, performed data collection, conducted bioinformatic data curation, conducted data analysis and produced visualizations for the manuscript. M.M, M.W., S.E.W. wrote the original draft of the manuscript with help from C.H.G., R.S., L.D.O.S., M.S.C., P.C., J.M.A.M., T.W., L.D. All authors reviewed and edited the manuscript. T.W. and L.D. conceived the project, provided supervision and acquired funding. All authors have read and agreed to the published version of the manuscript.

