## Supplementary files for "Maramycin, a cytotoxic isoquinolinequinone terpenoid produced through heterologous expression of a bifunctional indole prenyltransferase /tryptophan indole-lyase in *S. albidoflavus*"

**Table of Contents**

1) **Supplementary Figures** **S2-S10**

**Figure S1:** HPLC Traces of *S. albidoflavus* J1074 of WT and ΦC31::*pAzd* strains.

**Figure S2: (a)** UV and **(b)** HRMS spectra of maramycin (**1**)

**Figure S3:** 13C NMR Spectrum of Maramycin in DMSO-*d*6

**Figure S4:** Edited HSQC Spectrum of Maramycin

**Figure S5:** 1H NMR Spectrum of Maramycin in DMSO-*d*6

**Figure S6:** N-HMBC Spectrum of Maramycin

**Figure S7:** DQF-COSY Spectrum of Maramycin

**Figure S8:** HMBC Spectrum of Maramycin

**Figure S9:** Maramycin's Effect on Prostate Cancer Cell Proliferation

**Figure S10:** Plasmid Map of pRM4e-Mara1

2) **Supplementary Tables** **S11**

**Table S1:** NMR Data of Maramycin in DMSO-d6

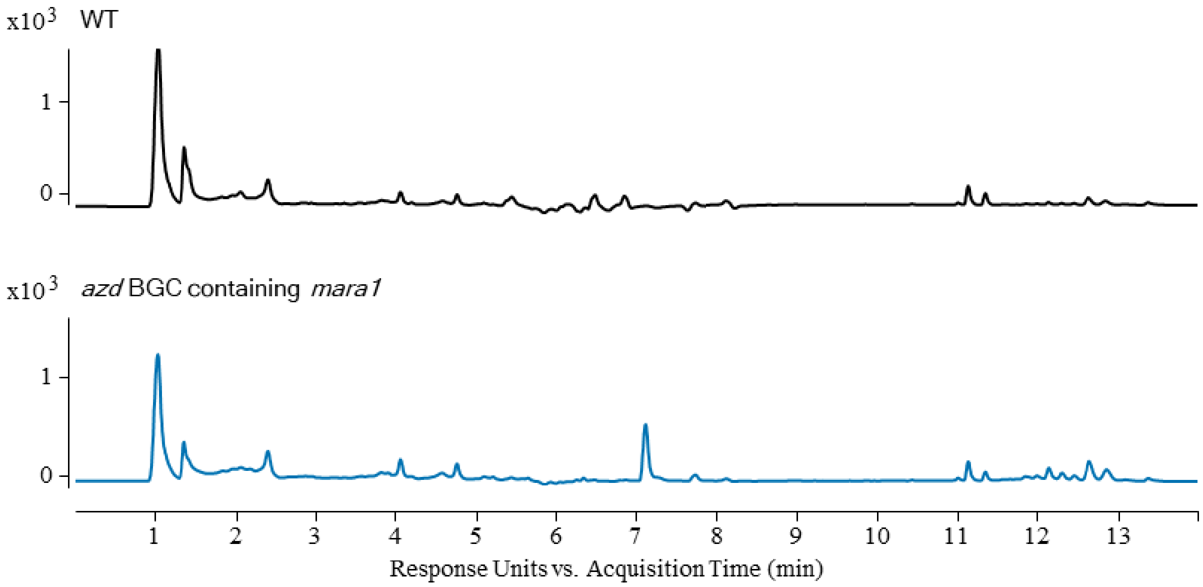


**Figure S1.** HPLC traces (280 nm) of LC-MS extracts prepared from *S. albidoflavus* J1074 (WT) and *S. albidoflavus* J1074 ΦC31::pAzd (*azd* BGC containing *mara1*).

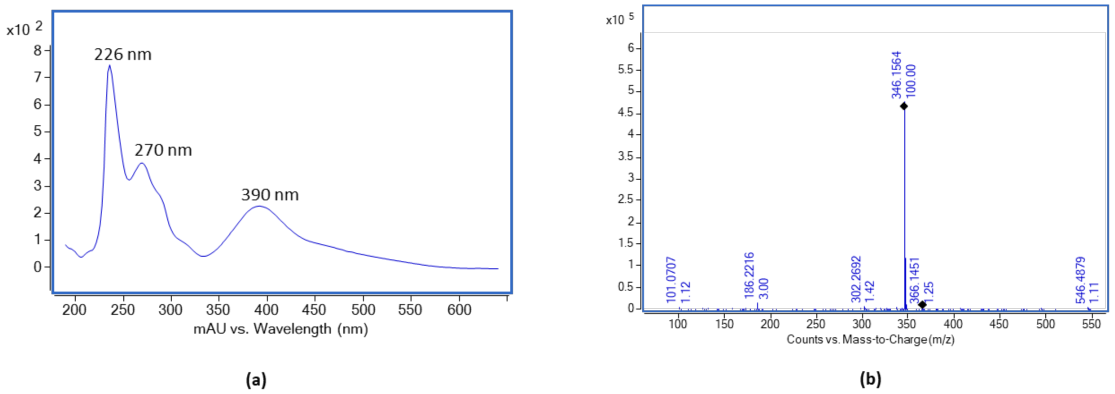


**Figure S2. (a)** UV and **(b)** HRMS spectra of maramycin (**1**)

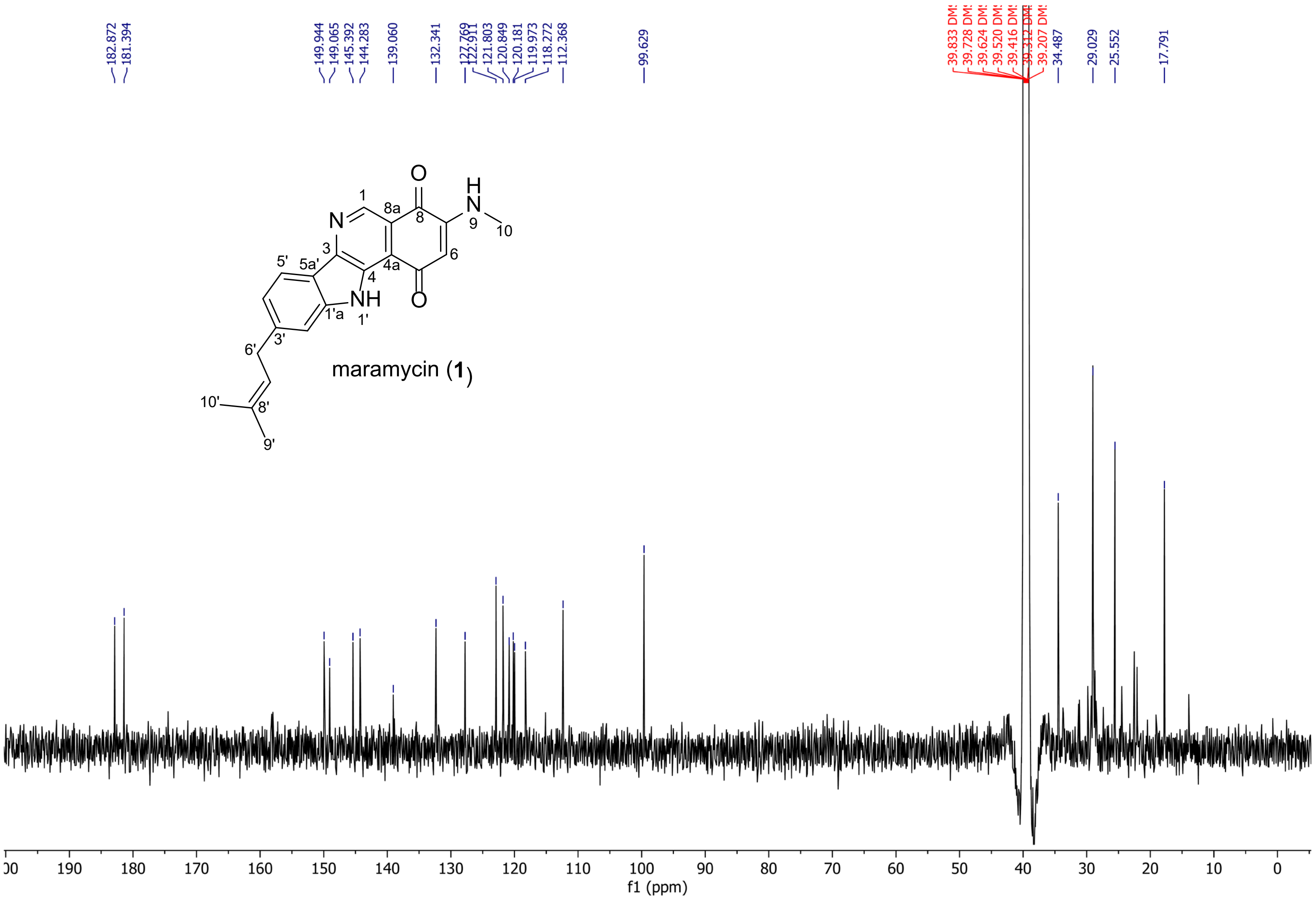


599

600 **Figure S3.** $^{13}C$ NMR (200 MHz) spectrum of maramycin (**1**) in DMSO-*d6*.

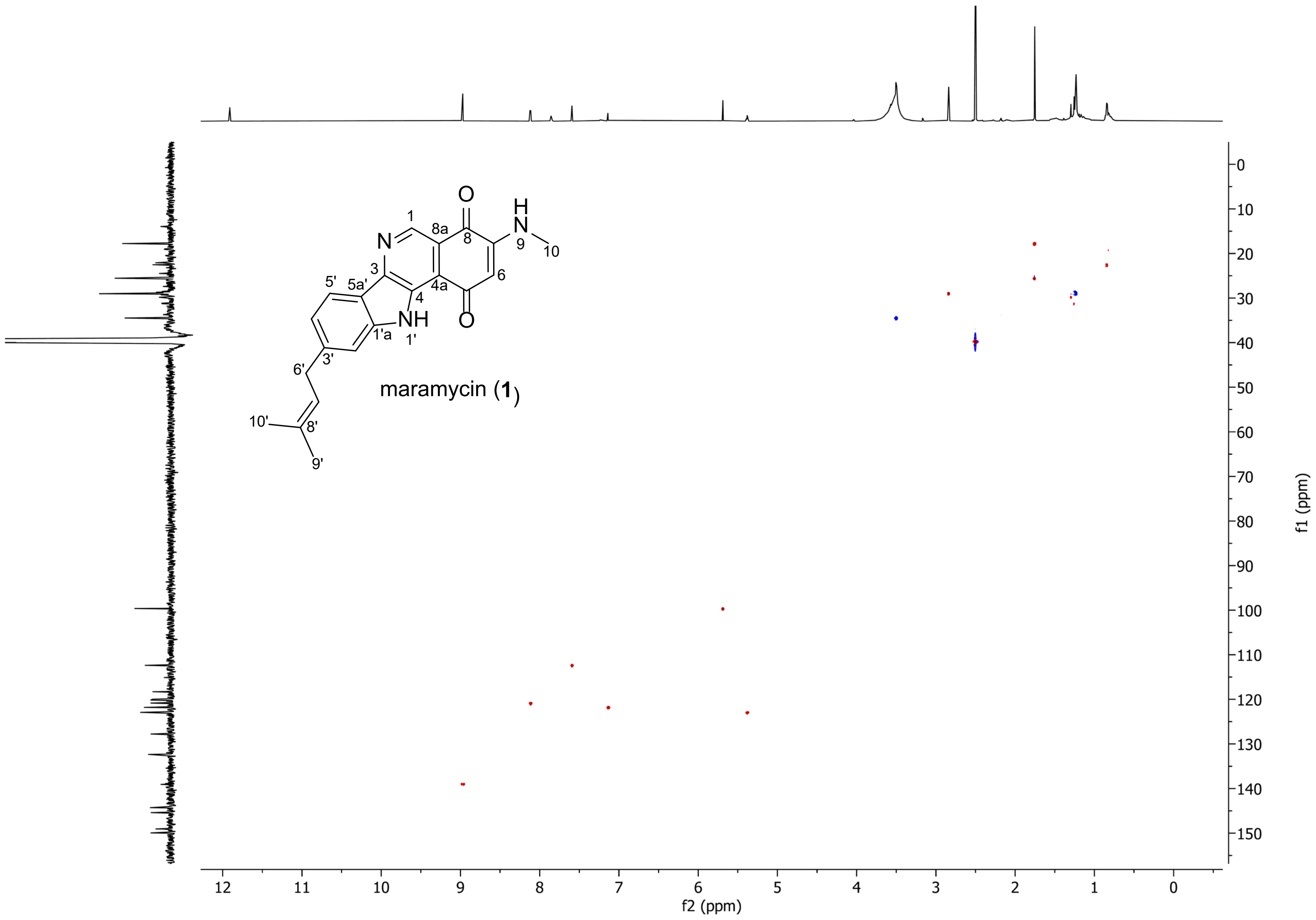


601

602 **Figure S4.** Edited HSQC spectrum of maramycin (**1**) in DMSO-*d*6.

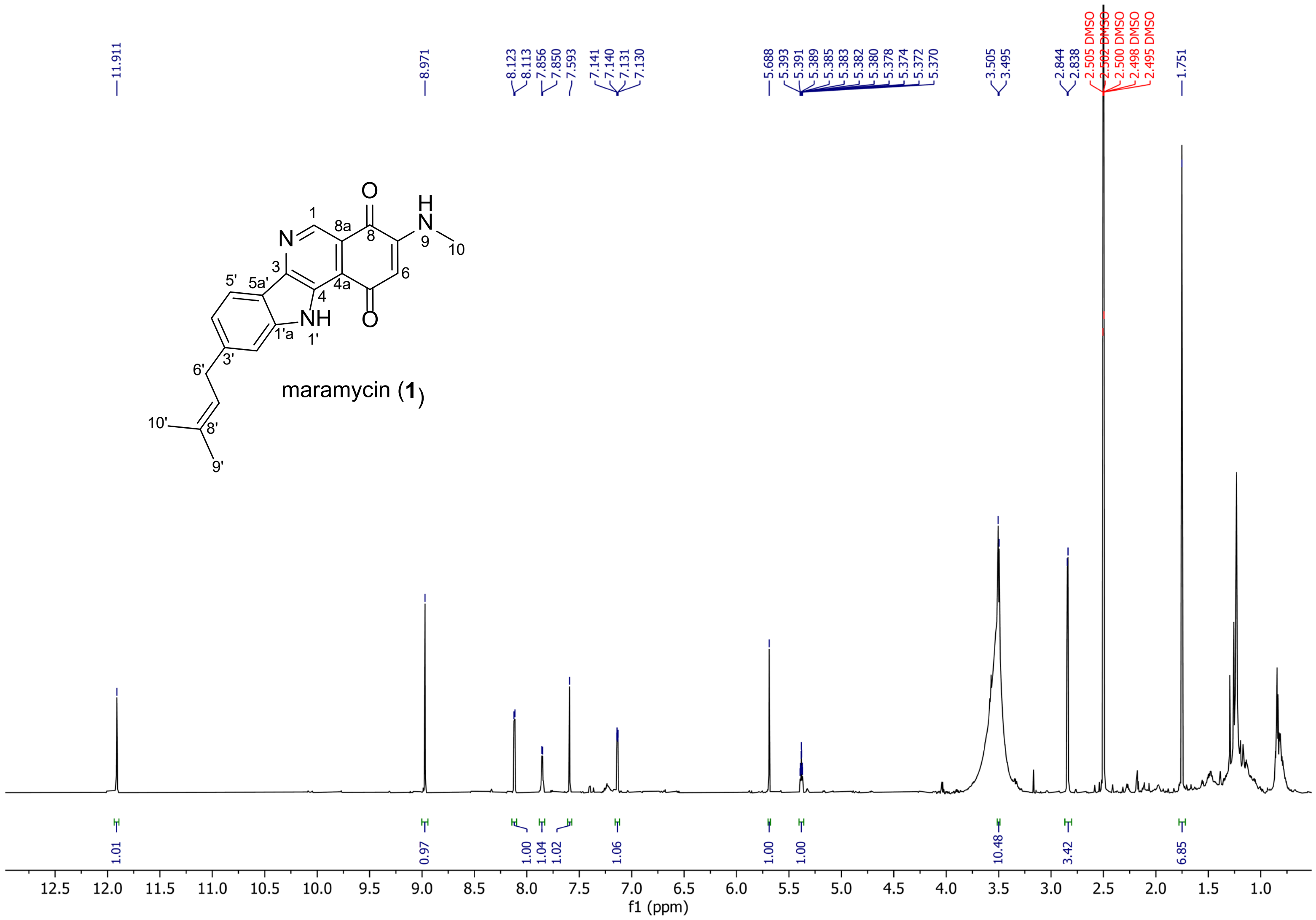


**Figure S5.** $^1$H NMR (800 MHz) spectrum of maramycin (**1**) in DMSO-*d6*.

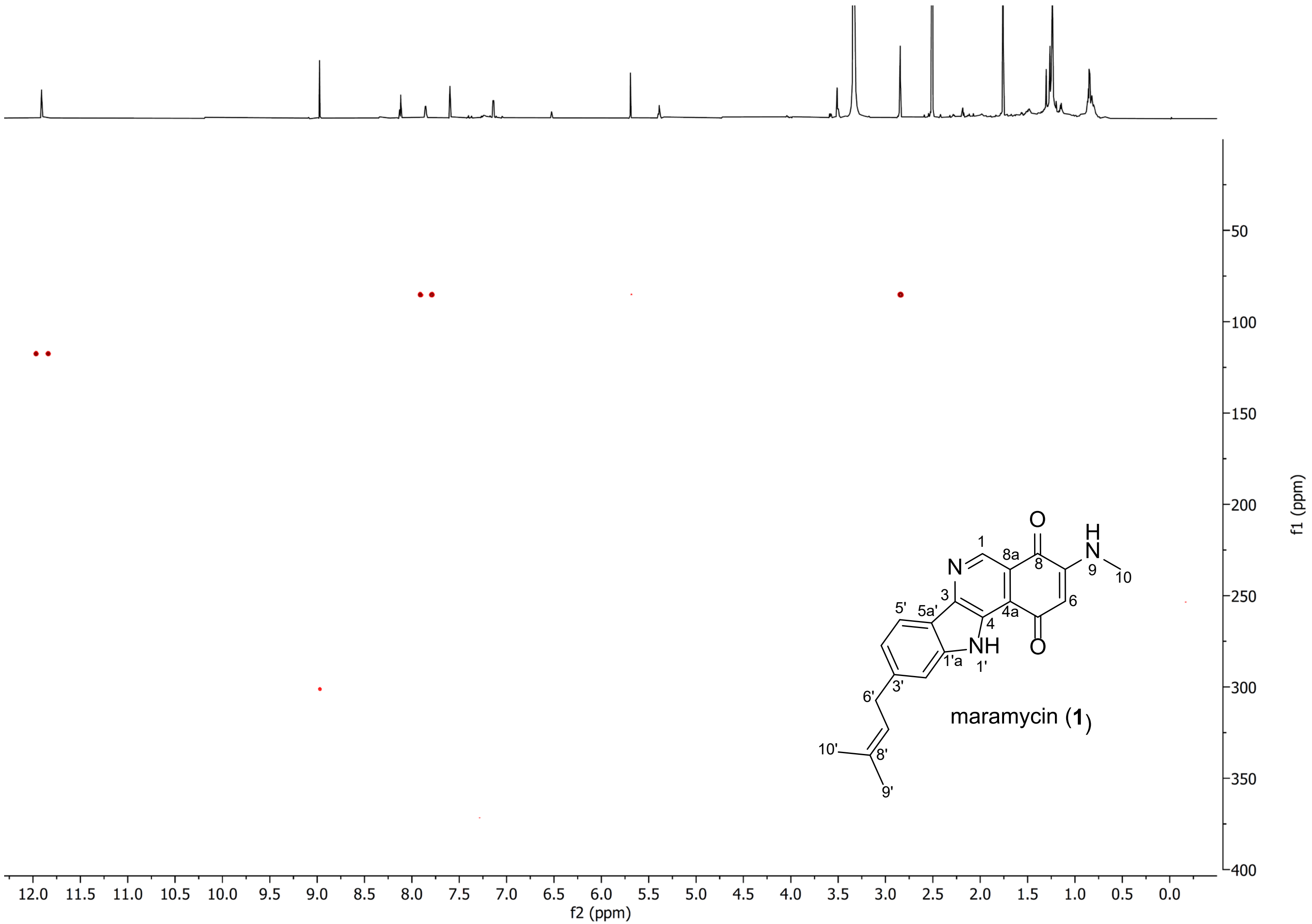


**Figure S6.** N-HMBC spectrum of maramycin (**1**) in DMSO-*d*6

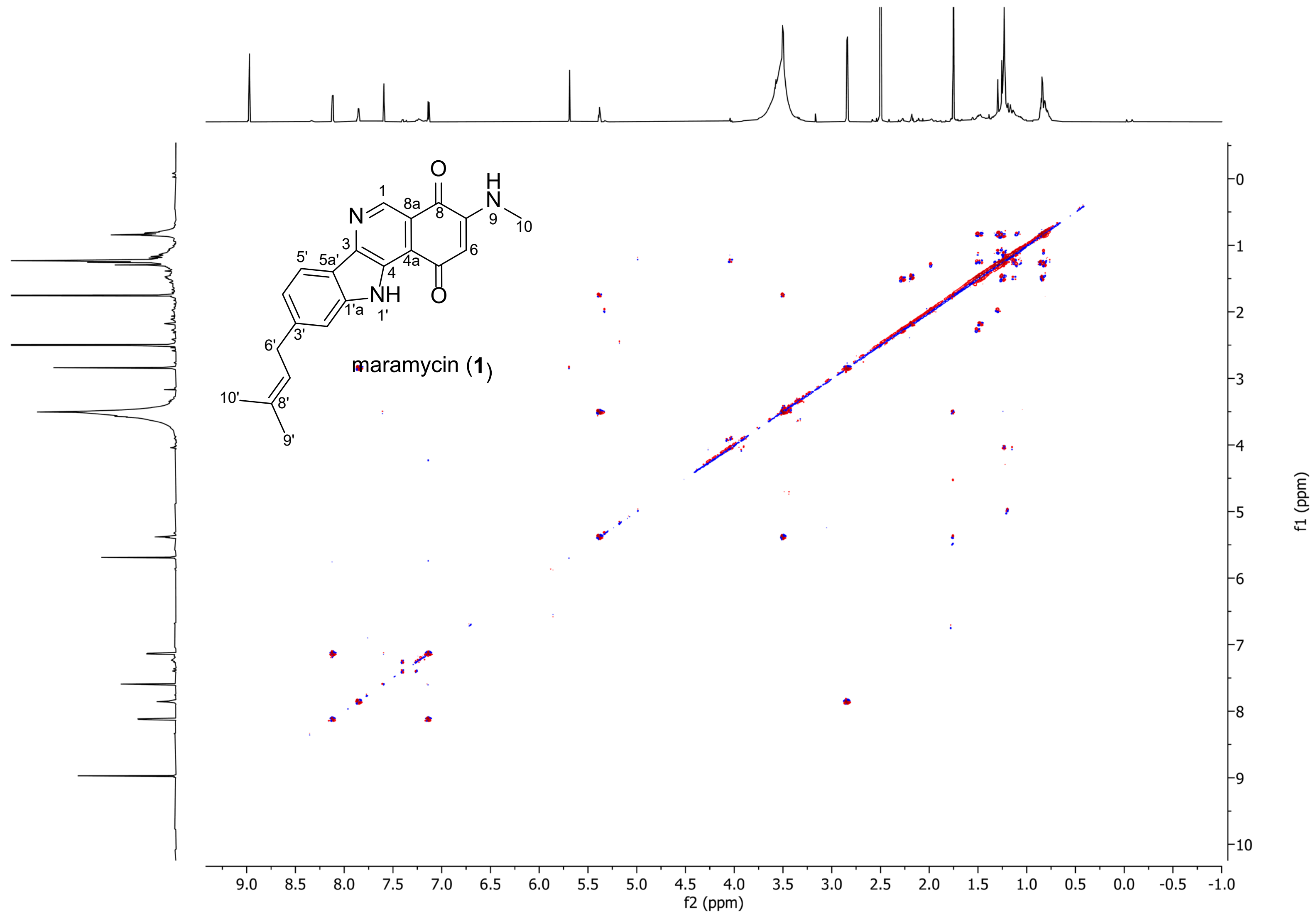


**Figure S7.** DQF-COSY spectrum of maramycin (**1**) in DMSO-*d6*.

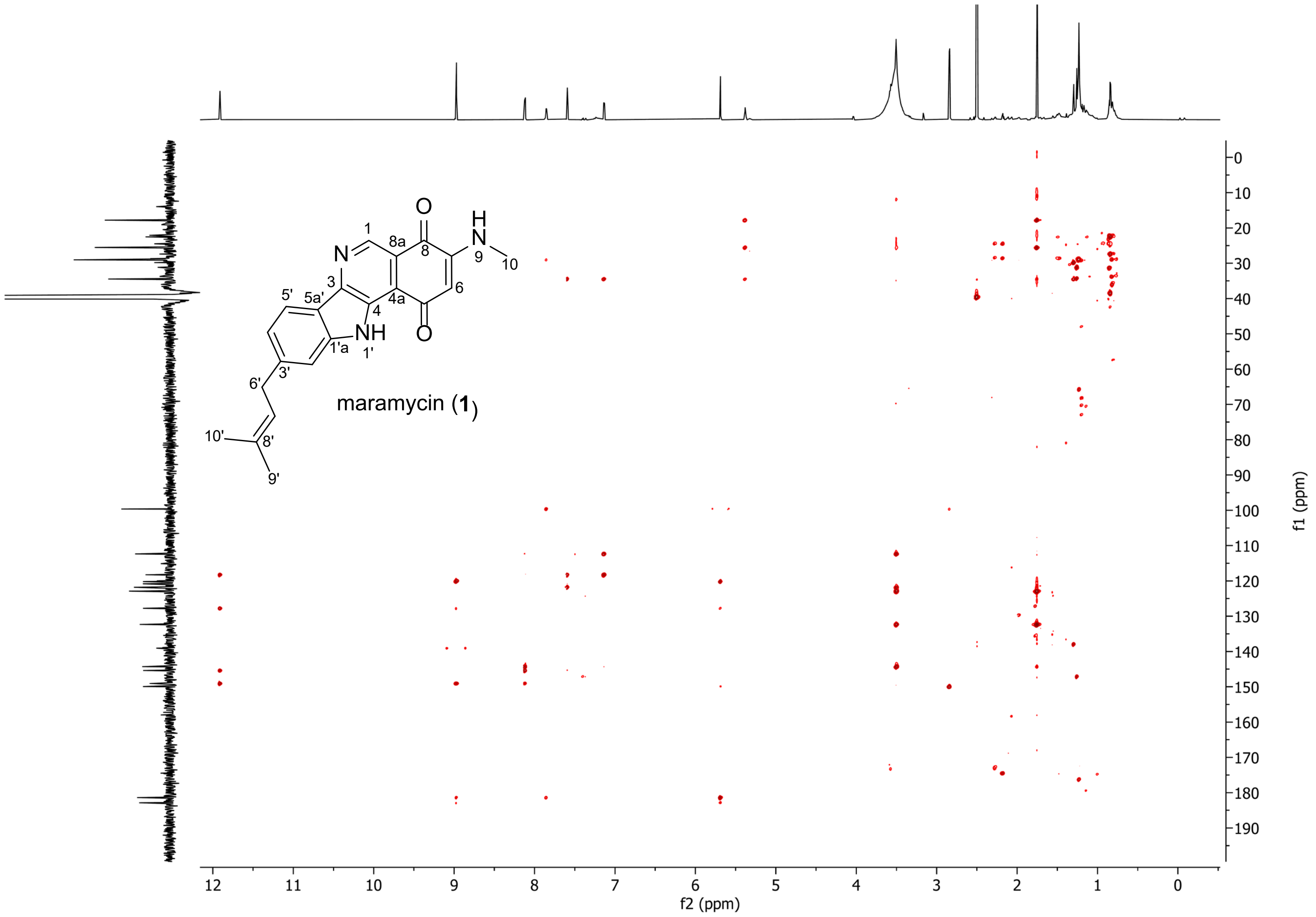


**Figure S8.** HMBC spectrum of maramycin (**1**) in DMSO-*d*6.

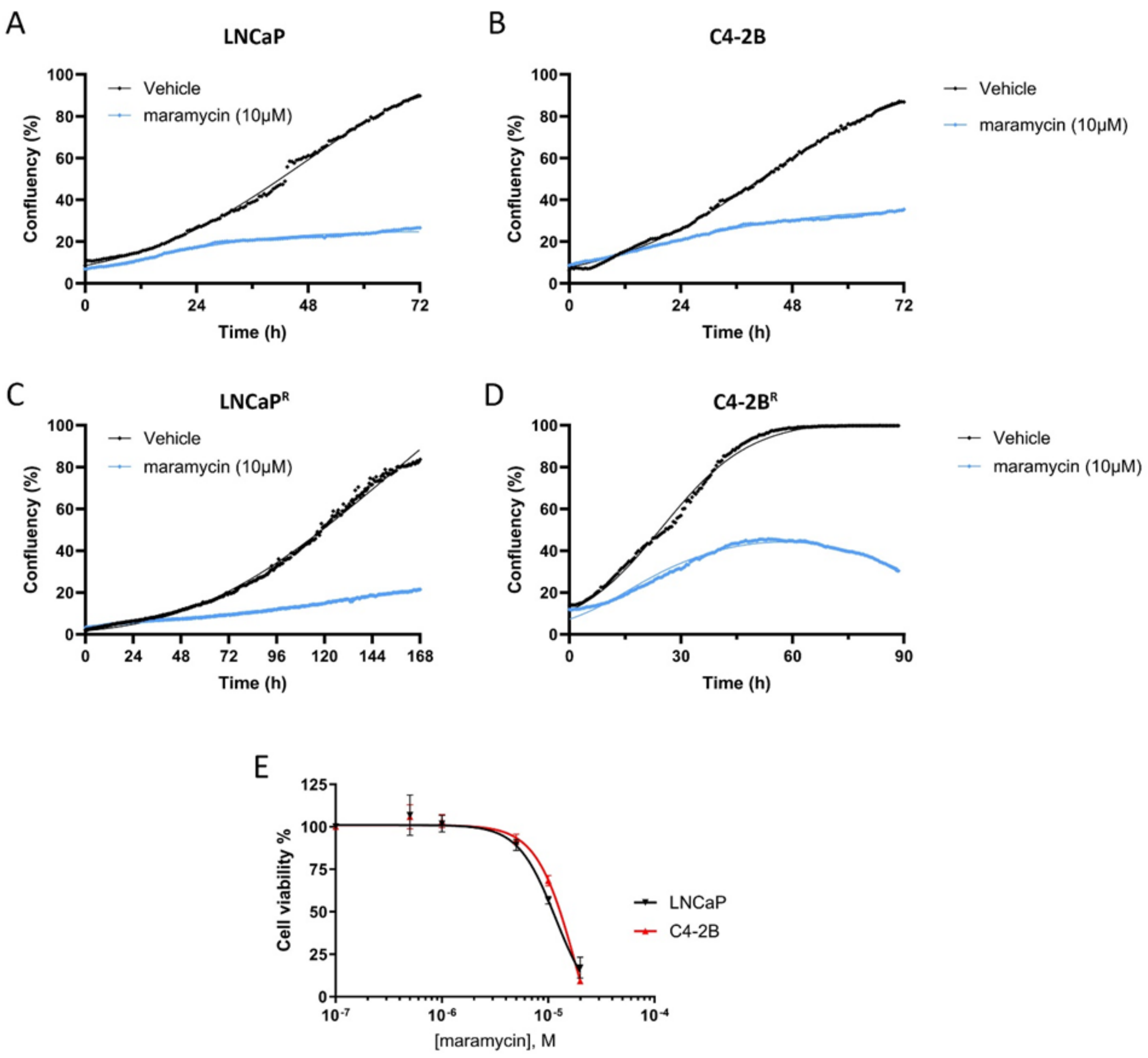


**Figure S9.** Maramycin affects cell proliferation and viability. Label-free confluence measurements of (A) LNCaP, (B) C4-2B, (C) LNCaP$^{R}$, and (D) C4-2B$^{R}$ prostate cancer cells, grown in the presence of 10µM maramycin or vehicle, respectively, show growth-inhibitory effects. (E) Determination of $IC_{50}$ values of maramycin for LNCaP and C4-2B cells, respectively. Plotted values were normalized to the untreated control.

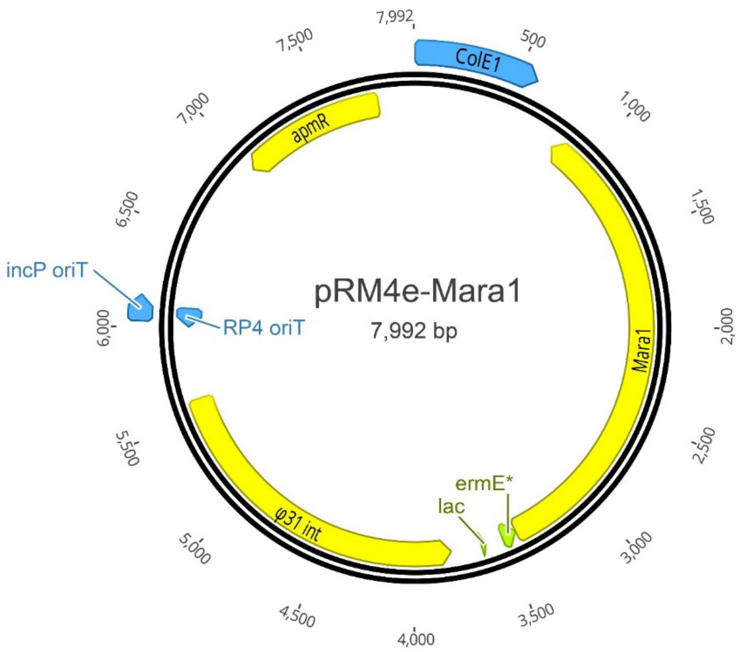


**Figure S10.** Plasmid map of pRM4e-Mara1. Generated using Geneious v.2023.2, Biomatters.

**Table S1**. $^{1}H$ (800 MHz), $^{13}C$ (200 MHz) and $^{15}N$ (80 MHz) NMR data of **1** in DMSO-$d_6$

| Position | $\delta_C$, type | $\delta_H$ ($J$ in Hz) | $\delta_N$, type[a] | HMBC[b] |
|---|---|---|---|---|
| 1 | 139.1, CH | 8.97, s | - | 3, 4a, 8 (2) |
| 2 | - | - | 301.1, N | |
| 3 | 149.1, C | - | - | |
| 4 | 127.8, C | - | - | |
| 4a | 120.2, C | - | - | |
| 5 | 182.9, C | - | - | |
| 6 | 99.6, CH | 5.69, s | - | 4a, 5, 8 |
| 7 | 149.9, C | - | - | |
| 8 | 181.4, C | - | - | |
| 8a | 120.0, C | - | - | |
| 9 | - | 7.85, q (5.1) | 85.3, NH | 6, 8, 10 (9) |
| 10 | 29.0, $CH_3$ | 2.84, d (5.1) | - | 7 (9) |
| 1′ | - | 11.91, s | 117.6, NH | 3,4,1’a,5’a (1’) |
| 1′a | 145.4, C | - | - | |
| 2′ | 112.4, CH | 7.59, s | - | 4’, 5’a, 6’ |
| 3′ | 144.3, C | - | - | |
| 4′ | 121.8, CH | 7.14, dd (8.1, 1.4) | - | 2’, 5’a, 6’ |
| 5′ | 120.8, CH | 8.12, d (8.1) | - | 3, 3’, 1’a |
| 5′a | 118.3, C | - | - | |
| 6′ | 34.5, $CH_2$ | 3.50, d (7.8) | - | 2’, 3’, 4’, 7’, 8’ |
| 7′ | 122.9, CH | 5.38, m | - | 6’, 9’, 10’ |
| 8′ | 132.3, C | - | - | |
| 9′ | 25.6, $CH_3$ | 1.75 (3H), s | - | 7’, 8’, 10’ |
| 10′ | 17.8, $CH_3$ | 1.75 (3H), s | - | 7’, 8’, 9’ |

[a]Assigned by $^{1}H$-$^{15}N$ HMBC correlation. [b] Number given in (x) is $^{1}H$-$^{15}N$ correlation.

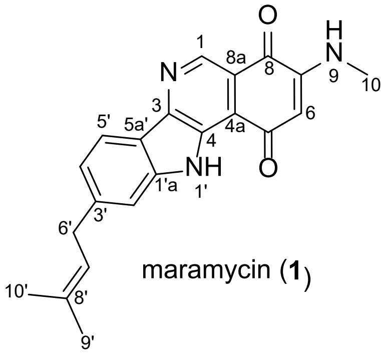


maramycin (**1**)
